# Bioinspired Virus-Like Porous Silica Amplify Lipid-Mediated mRNA Delivery

**DOI:** 10.64898/2026.05.02.722380

**Authors:** Silja Saarela, Kirsti Härkönen, Milla-Iida Laari, Minna Sivonen, Tomas Strandin, Jussi Hepojoki, Einari Niskanen, Vesa-Pekka Lehto, Wujun Xu

## Abstract

Lipid nanoparticles (LNPs) have demonstrated strong potential in COVID-19 mRNA vaccines nevertheless they still face the challenges in low mRNA delivery efficacy. Virus-like porous silica (VLPSi) nanoparticles (NPs) represent a promising biomimetic delivery platform because their spiked morphology may enhance cellular internalization and promote endosomal membrane disruption. However, the application of VLPSi for mRNA has been rarely explored. In this study, hybrid lipid–VLPSi NPs were developed by combining VLPSi with either lipoplexes (LPs) or LNPs. The effects of lipid types, mass ratio of different compositions, and amine modifications of VLPSi on mRNA delivery were studied. The results demonstrated that both LP and LNP could be successfully integrated with VLPSi to form hybrid delivery systems for mRNA transfection. VLPSi could significantly enhance mRNA delivery of both LPs and LNPs due to improved cellular uptake, structural stabilization of the mRNA complex, and enhanced endosomal escape mediated by the rigid virus-like surface architecture. Among the tested lipid formulations, the ionizable lipid ALC-0315 and helper lipid DOPE with mass ratio of 5:3 was the most effective lipid composition to be integrated with VLPSi, showing the highest mRNA delivery performance. In addition, amino modification of VLPSi was found to be a critical factor for efficient mRNA delivery. Hybrid LNPs containing amino-modified VLPSi showed significantly higher transfection efficiency than those containing unmodified VLPSi. Notably, amino-modified LNP–VLPSi achieved up to fivefold higher gene expression than conventional LNPs. Overall, this study establishes VLPSi as an efficient platform for amplifying lipid-mediated mRNA delivery. Owing to its straightforward integration into widely used LNP systems, VLPSi offers an adaptable and effective strategy for advancing next-generation mRNA therapeutics.

## 1 Introduction

The COVID-19 pandemic was a turning point in the development of vaccines [1]. Unlike previous vaccines, a new class of severe acute respiratory syndrome coronavirus 2 (SARS-CoV-2) vaccines contain neither viral protein nor inactivated pathogens to trigger immunity [2]. Instead, these vaccines consist of synthetic messenger-RNA (mRNA), encoding SARS-CoV-2 spike glycoprotein [3]. Since then, mRNA vaccines have gained more attention, offering advantages, such as carrying multiple antigens and rapid design and synthesis at low production costs [4]. Moreover, the entering of mRNA vaccine in cell cytoplasm is enough to enable antigen expression, resulting no risk for genomic alterations or mutations [5]. Thus, mRNA vaccines provide high safety and specificity [6]. However, mRNA needs a carrier to protect it from the rapid degradation in the extracellular environment [7]. Moreover, a carrier is needed to pass electro statical physiological barriers and finally, release mRNA from endo-lysosomal system to cell cytoplasm [8].

Lipid nanocarriers, such as lipoplexes (LPs) and lipid nanoparticles (LNPs), have proven their ability as an functional mRNA delivery system [9]. Common LP formulation consist of a cationic or ionizable lipid and a helper lipid [10]. In addition to these, LNPs typically contain cholesterol and polyethylene glycol-linked lipid (PEG-lipid), enhancing their structure and stability compared to LPs [11,12]. In both lipid nanocarriers, cationic or ionizable lipid plays a crucial role in mRNA encapsulation [13]. Even if positively charged cationic lipids can increase mRNA loading capacity, they might induce undesirable cytotoxicity [14]. On the other hand, pH dependent ionizable lipids, such as ALC-0315, have garnered interest, being neutral at physiological pH, but positively charged under acidic conditions [2]. Thus, ionizable lipids provide lower cytotoxicity and enhanced biocompatibility while their destabilizing non-bilayer structures at low pH enable efficient endosomal escape by promoting membrane disruption [2,15]. Beside ionizable lipids, uncharged helper lipids are primary component of lipid nanocarriers [16]. The choice of helper lipid can affect the properties of the nanocarrier, such as stability, mRNA encapsulation capacity, or endosomal escape [16,17]. For example, among the commonly used helper lipids, 1,2-distearoyl-sn-glycero-3-phosphocholine (DSPC) is known to improve stability whereas 1,2-dioleoyl-sn-glycero-3-phosphoethanolamine (DOPE) enhances endosomal escape [18]. However, despite the success of lipid nanocarriers in mRNA vaccines, LNPs can deliver less than 2% of the administered mRNA to cells due to their inefficient endosomal escape [19]. Thus, improved mRNA nanocarriers are needed.

Inspired by viral structures, several different virus-like nanoparticles have been developed for biomedical applications [20]. Among these, mesoporous silica nanoparticles (MSNs) have attracted considerable attention as nanocarriers due to their large surface area, high biocompatibility and structural stability. Due to their tuneable morphology, rigid virus-like spikes are easily introduced on the MSN surfaces to obtain virus-like mesoporous silica nanoparticles (VLPSi) [20–22]. Beside the advantageous properties of MSNs, the viral morphology of VLPSi promotes the cell membrane binding affinity and cell internalization [23]. Similarly, the spikes of VLPSi can penetrate through endosomal membrane. This aids endosomal escape, being especially beneficial for appropriate mRNA delivery [24]. However, mRNA molecule is too large to be loaded into mesopores of VLPSi [25,26], but the surface spikes on VLPSi might provide protective microenvironment for mRNA [24]. Still, to bound the negatively charged mRNA into negatively charged VLPSi via electrostatic interaction, positive linker, such as ionizable or cationic lipid, between mRNA and VLPSi is needed [27]. However, the application of combined LNPs and VLPSi for mRNA delivery has not been studied.

In the present study, we developed hybrid lipid-VLPSi to act as an efficient mRNA delivery system. Hence, VLPSi were combined with both LPs and LNPs to evaluate their mRNA transfection efficiency, using mCherry-encoding mRNA. First, LPs were composed of ALC-0315 and 1,2-Dioleoyl-sn-glycero-3-phosphocholine (DOPC) to select the optimal mass ratio between ionizable and helper lipid. Moreover, different mass ratios between LPs and VLPSi were tested. Further, different cationic lipids and helper lipids were used to investigate the optimal LP formulation. Finally, the optimized lipid composition was used to integrate LNP with VLPSi with and without amino-modification for mRNA transfection.

## 2 Materials and methods

### 2.1 In vitro transcription of mRNA

mCherry mRNA was transcribed from mCherry carrying plasmid DNA (pDNA) provided by Jussi Hepojoki”s laboratory. pDNA was used as a template for polymerase chain reaction (PCR) to amplify the mCherry gene with T7 promoter based on the method previously published by Ravi Ojha et al. (2025) [28]. Briefly, Q5 High-Fidelity 2x master mix (New England BioLabs, USA) was used in the PCR protocol described more detailed by Ravi Ojha et al. (2025) [28]. Amplified DNA was purified using NucleoSpin Gel and PCR Clean-up kit (Macherey-Nagel, Germany) following the manufacturer”s instructions. Further, HiScribe T7 ARCA mRNA Kit (New England BioLabs, USA) was utilized in *in vitro* transcription and purification of mCherry mRNA based on the kit protocol when amplified DNA was used as a template. Subsequently, Poly-A tailing of mRNA was conducted. Finally, the concentration of purified mCherry mRNA in RNase-free water was measured using NanoDrop One UV-Vis Spectrophotometer (Thermo Fisher Scientific, USA). The final product was stored at -80 °C.

To confirm the production and purity of DNA and mRNA and Poly-A tailing of mRNA, the products were run by agarose gel electrophoresis (2% and 1.5% agarose gels containing 0.01% of GelRed, respectively). In the preparation of DNA samples, DNA Loading Dye, Tritrack DNA loading dye, and GeneRuler 1 kb DNA Ladder (Thermo Fisher Scientific, USA) were used. Respectively, RiboRuler Low Range RNA ladder and RNA Loading Dye (Thermo Fisher Scientific, USA) were used to prepare RNA samples following the manufacturer”s protocol. The gels were run with 100 V until the bands were clearly visible. Afterwards, the gels were imaged using ChemiDOC MP Imaging System (Bio-Rad, USA) (**Figure S1**).

### 2.2 Preparation of lipid reagents

#### 2.2.1 LPs

LPs were prepared using thin-film hydration method. First, the stock solutions of cationic lipids, 1,2-Dioleoyl-3-trimethylammonium-propane chloride (DOTAP, MedChemExpress, USA) and 1,2-di-O-octadecenyl-3-trimethylammonuim propane (DOTMA, Indagoo, Europe), ionizable lipid ALC-0315 (MedChemExpress, USA), and helper lipids, DOPE (MedChemExpress, USA) and DSPC (MedChemExpress, USA) were prepared in chloroform. The third helper lipid, DOPC (Avanti Research, USA) was originally in chloroform at the concentration of 25 mg/ml. To prepare ALC-0315:DOPC (1:1, 5:3, 2:1, 3:1, 4:1, and 5:1, w/w), ALC-0315:DOPE (5:3, w/w), and ALC-0315:DSPC (5:3, w/w), the desirable amounts of ALC-0315 and DOPC, DOPE or DSPC from the stocks were mixed in a glass vial. To verify the successful pipette mixing, a certain amount of extra chloroform was added so that the final volume of each lipid mixture was 100 µl. The chloroform was allowed to evaporate at room temperature (RT) turning the vial occasionally by hand to enhance evaporation and mixing. After evaporation, the formed LPs were recollected using 80% ethanol in MilliQ so that their final concentration was 1 mg/ml. Respectively, DOTAP:DOPE (1:2, 1:1, 2:1, w/w) and DOTMA:DOPE (1:2, 1:1, 2:1, w/w) were prepared. Finally, the LP stocks were stored at - 20 °C.

#### 2.2.2 Lipids for LNPs

Stock solutions of ALC-0315, DOPE, and PEG-lipid, 2-[(Polyethylene glycol)-2000]-N,N-ditetradecylacetamide (ALC-0159, MedChemExpress, USA), were prepared in ethanol at the concentrations of 1 mg/ml (ALC-0315 and DOPE) and 0.5 mg/ml (ALC-0159). Further, ALC-0315, DOPE, and ALC-0159 from the stocks were mixed with pipette in the mass ratio of 58.6 : 35.2 : 6.2 to be subsequently utilized in the preparation of LNPs for mRNA transfection.

### 2.3 Preparation of VLPSi

#### 2.3.1 VLPSi

The synthetization of VLPSi followed the previously published method of Wenxing Wang et al. (2017) [29]. The published method is based on epitaxial growth approach, in which cationic surfactant, cetyltrimethylammonium bromide (CTAB), is used as a template while tetraethyl orthosilicate (TEOS) and sodium hydroxide (NaOH) are acting as a silica source and a catalyst, respectively. Cyclohexane is used as an organic solvent to form the oil phase [29].

Beginning of the synthesis, 0.1 g of CTAB and 10 ml of MilliQ water were transferred in a flat-bottom flask, which was placed in a 60 °C oil bath with stirring. Gradually, 0.15 ml of 0.1 M NaOH was added into the solution, which was allowed to react for 2 h. Next, 20 ml of TEOS mixed in 80 ml of cyclohexane was added. The reaction was continued at 60 °C with stirring rate 350 rpm for 72 h, aiming VLPSi with 30 nm spike length. After reaction, VLPSi were washed with both MilliQ water and ethanol via centrifugation at 10 000 rpm for 10 min. To ensure the complete removal of CTAB template, VLPSi were calcinated at 600 °C for 4 h and finally stored in ethanol.

#### 2.3.2 Amino-modified VLPSi

To turn VLPSi surface charge from negative to positive, free NH_2_, N(CH_3_)_3_, or NH-NH-NH_2_ groups were added on the surface of VLPSi. First, 3 µl of pure (3-Aminopropyl)triethoxysilane (APTES, Acros Organics, Belgium), 6 µl of N-[3-(Trimethoxysilyl)propyl]-N,N,N-trimethylammonium chloride in 50% ethanol (Thermo Scientific, USA), or (3-Trimethoxysilylpropyl)diethylenetriamine (abcr GmbH, Germany) was mixed with 5 mg of VLPSi in 1.5 ml of ethanol. The mixtures were stirred at 65 °C for 1 h. After reaction, the obtained amino-modified VLPSi were washed three times with ethanol via centrifugation at 14 000 rpm for 10 min.

#### 2.3.3 FITC-NH_2_-VLPSi-COOH

To study cell uptake, NH_2_-VLPSi were labelled with fluorescein isothiocyanite (FITC, Alfa Chemical, China). Simultaneously, to turn positive surface charge of NPs to negative, succinic anhydride was used to cap free NH_2_ groups by forming COOH groups on the NP surfaces. Thus, 0.5 mg of FITC, 5 mg of succinic anhydride, and 5 mg of NH_2_-VLPSi were mixed in 1 ml of ethanol. The mixture was rotated overnight at RT in the dark. Next day, obtained FITC-NH_2_-VLPSi-COOH were washed three times in ethanol by centrifuging at 14 000 rpm for 10 min.

### 2.4 Physicochemical characterizations

Transmission electron microscopy (TEM, Jeol JEM-2100F, Japan) was used to study the morphology and size of VLPSi. For sample preparation, 10 µl of 0.1 mg/ml VLPSi suspended in ethanol was placed on an EMR carbon support film on copper grid (400 square mesh, Micro to Nano, Netherlands). The grid was dried at RT until ethanol was evaporated.

Zeta potentials of both VLPSi with and without amino-modifications and transfection complexes was measured using Zetasizer Nano ZS (Malvern, UK). For the measurement, NPs and complexes were suspended in MilliQ water.

### 2.5 Cell uptake of VLPSi and LP-VLPSi

HEK-293T cells were seeded into 8-well plate (Ibidi, Germany) at the density of 20 000 cells/well and cultured at 37 °C overnight. Then, the cells were treated with 10 µg/ml of ALC-0315:DOPE (5:3, w/w)- and FITC-NH_2_-VLPSi-COOH for 1 and 4 h. After treatment, cells were washed once with PBS to remove free NPs. Cell nuclei were stained with 10 µg/ml of Hoechst 33342 (Sigma-Aldrich, USA). The cells were incubated for 10 min and imagined with Zeiss LSM 800 Airyscan (Zeiss, Germany). However, to study the cell uptake at 24 h time point, cell incubation in culture medium was continued after the removal of 4 h incubated NPs, followed by nuclei staining and imaging respectively.

### 2.6 mRNA transfection in vitro

#### 2.6.1 Cell culture

To study mRNA transfection, cells were seeded in 24-well plates (Greiner Bio-One Polystyrene 24-well Cell Culture Multiwell Plate, Germany) according to **Table 1** and cultured at 37 °C for 24 h. The culture mediums contained 10% of fetal bovine serum (FBS, Gibco™, Thermo Fisher Scientific, USA) and 1% of penicillin-streptomycin (P/S, Gibco™, Thermo Fisher Scientific, USA). For HEK-293T and SK-N-SH cells, well plates were coated with Poly-L-Lysine to enhance the cell adhesion. To avoid the macromolecules of FBS to disturb the internalization of transfection complexes, culture medium was replaced with fresh serum-free medium before transfection. The amount of fresh medium per well was optimized to be 200 µl and 350 µl for VLPSi and lipid reagents, respectively.

**Table 1.** Cell lines used to study mRNA transfection *in vitro*.

| Cell line | Origin | Culture medium | Cells/well |
| --- | --- | --- | --- |
| Human embryonic kidney 293T (HEK-293T) | Human | DMEM with 4.5 g/L glucose, with L-Glutamine, without sodium pyruvate, 10 % FBS, 1% penicillin-streptomycin | 60 000 |
| Human triple negative breast cancer (MDA-MB-231) | Human | RPMI 1640 with L-glutamine, 10 % FBS, 1% penicillin-streptomycin | 100 000 |
| Human neuroblastoma (SK-N-SH) | Human | DMEM with 4.5 g/L glucose, with L-Glutamine, without sodium pyruvate, 10 % FBS, 1% penicillin-streptomycin | 100 000 |
| Human monocytic leukemia (THP-1) | Human | RPMI 1640 with L-glutamine, 10 % FBS, 1% penicillin-streptomycin | 100 000 |
| Macrophage-like (RAW264.7) | Mouse | DMEM with 4.5 g/L glucose, with L-Glutamine, without sodium pyruvate, 10 % FBS, 1% penicillin-streptomycin | 80 000 |

#### 2.6.2 mRNA transfection with LPs

To study mRNA transfection with LPs, Lipofectamine™ 2000 (Invitrogen™, USA) was used as a positive control according to the manufacturer”s instructions. The mRNA to Lipofectamine mass ratio was 1:4. Furter, LPs were dissolved in Opti-MEM medium (Gibco™, Thermo Fisher Scientific, USA) at predetermined concentrations. Subsequently, mCherry mRNA in citrate buffer (pH 4.0) was combined with LP solutions. Immediately after, the solutions were briefly mixed with vortex (∼10 s) and incubated at RT for 15 min, allowing the formation of transfection complexes. Finally, 30 µl of complexes was distributed to each well so that the mass of mRNA per well was 150 ng, determining mRNA to LPs mass ratio to be 1:8. The complexes were incubated with cells for 4 h. After incubation, medium was replaced with 500 µl of fresh culture medium (10% FBS).

#### 2.6.3 mRNA transfection with LP-VLPSi

As in the section 2.6.2, LPs and mCherry mRNA were respectively dissolved in Opti-MEM and citrate buffer (pH 4.0) and combined via rapid vortex mixing. Now, VLPSi were washed with RNase-free water and redispersed in Opti-MEM at a concentration of 1 mg/ml. Subsequently, VLPSi were combined with mRNA-LP solutions at predetermined mass ratios between, mRNA, LPs, and VLPSi: ALC-0315:DOPC (1:4:75, 1:8:75, 1:12:75, 1:16:75, 1:24:75, 1:8:10, 1:8:25, 1:8:50, 1:8:150, and 1:8:225), and ALC-0315:DSPC, ALC-0315:DOPE, DOTAP:DOPE, and DOTMA:DOPE (1:8:25). Further, 500 µl of serum-free culture medium was added to each solution, which were again mixed with vortex and incubated at RT for 15 min. Finally, 180 µl of complexes was distributed into each well. The mRNA amount per well was 150 ng. After 4 h incubation, free complexes were removed by replacing medium with 500 µl of fresh culture medium (10% FBS) per well.

#### 2.6.4 mRNA transfection with LNPs

To form LNPs, lipids from stock solution prepared as in the section 2.2.2 were combined with mCherry mRNA in citrate buffer (pH 4.0) by rapidly mixing up and down with pipette. The volume ratio between mRNA (aqueous phase) and lipids (organic phase) was 3:1. After mixing, LNPs were allowed to form by incubating the solution at RT for 15 min. Then, LNPs were dissolved in Opti-MEM, determining the final volume of 100 µl. Further, 30 µl of LNP solution was distributed in each well so that the mRNA amount per well was 150 ng. The mRNA to lipid mass ratio was 1:8. LNPs were incubated with cells for 4 h before replacing medium with 500 µl of fresh culture medium (10% FBS) per well.

#### 2.6.5 mRNA transfection with LNP-VLPSi

To form LNP-VLPSi with pipette mixing method, VLPSi or amino-modified VLPSi in ethanol were mixed with lipid solution prepared as described in the section 2.2.2, forming an organic phase. Now, mCherry mRNA in citrate buffer (pH 4.0) was combined with formed organic phase by rapid pipette mixing in the volume ratio of 3:1. Then, the solution was incubated at RT for 15 min before adding 49.6 µl of Opti-MEM and 500 µl of serum-free medium, determining the final volume of 600 µl. Formed LNP-VLPSi were briefly mixed with vortex and distributed into wells (180 µl/well). The mass ratio between mRNA, LNPs, and VLPSi was 1:8:25 while mRNA amount per well was 150 ng. After 4 h incubation, free complexes were removed by replacing medium with 500 µl of fresh culture medium (10% FBS) per well.

#### 2.6.6 Flow cytometry

After 24 h from mRNA transfection, the cells were detached with 0.25% of trypsin-EDTA (Gibco™, Thermo Fisher Scientific, USA) and washed with 1 ml of cold flow buffer (0.02 % of NaN3, 2 mM of EDTA, and 2 % of FBS in PBS) at 400 rfc for 8 min. After washing, the supernatant was removed and the cells were resuspended into 100 µl of cold flow buffer. The mCherry mRNA expression and median fluorescence intensity (MFI) of the cells was obtained with CytoFLEX S (Beckman Coulter, USA) and further analysed using CytExpert v2 (Beckman Coulter, USA) and FlowJo v10 (BD Life Sciences, TreeStar, USA). The flow cytometry gating strategy for the analysis of mCherry positive cells is shown in **Figure S2**.

### 2.7 Statistical analysis

GraphPad Prism 10 software (Dotmatics, UK) was used for statistical data analysis. All the values are presented as mean ± standard deviation and compared using one-way and two-ways ANOVAs with Tukey”s test. P < 0.05 was used as a threshold for statistical significance.

## 3 Results

### 3.1 Physicochemical characterizations

VLPSi were synthetized based on epitaxial growth approach. After removal of surfactant template, obtained VLPSi were TEM imaged to evaluate their morphology and size. **Figure 1a** shows TEM image of VLPSi, confirming 30 nm spikes length and uniform size. The average diameter of VLPSi is 80.3 ± 5.1 nm (n = 20) measured using ImageJ.

**Figure 1.**
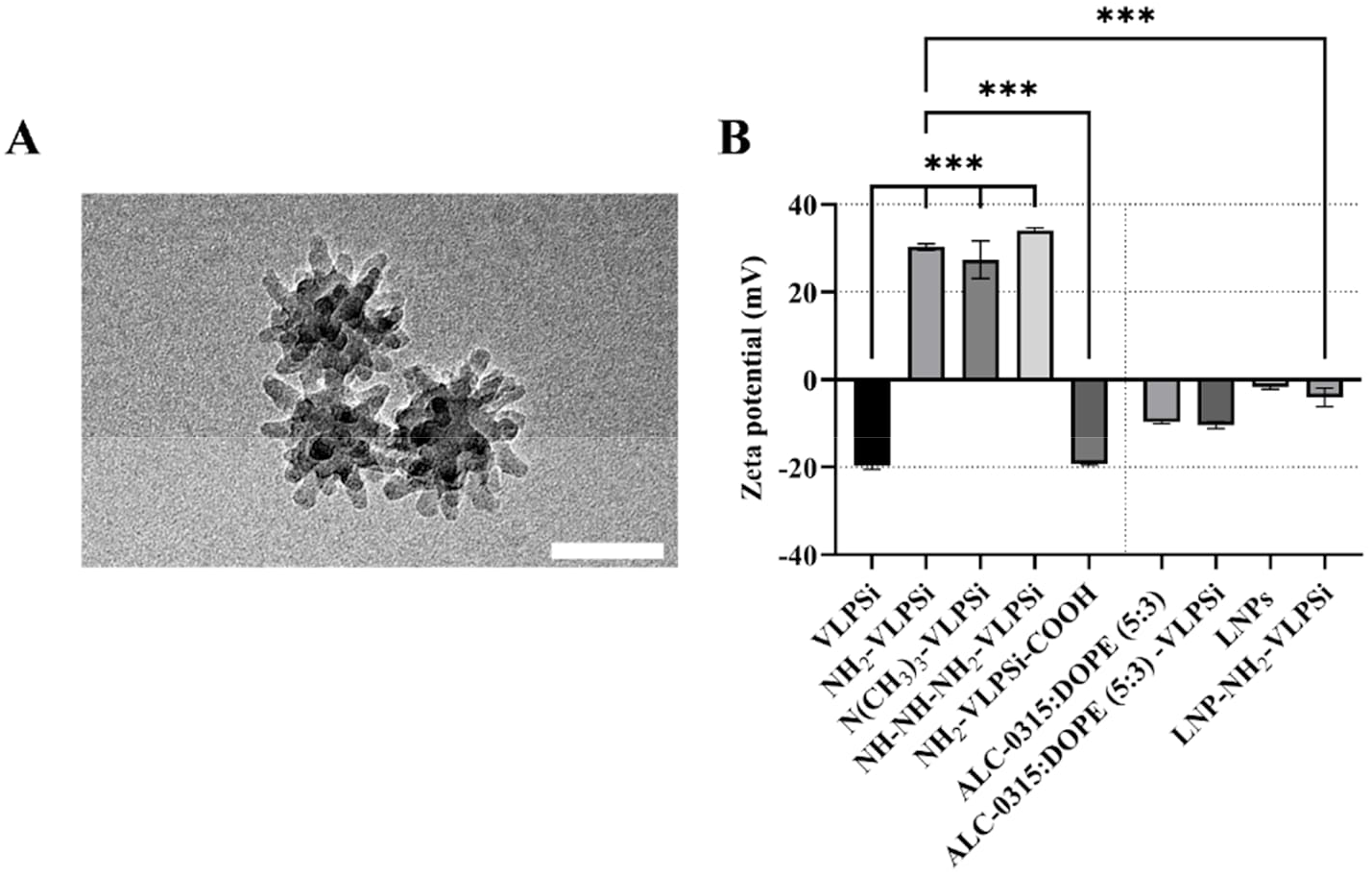
Physicochemical characterizations. **(A)** Transmission electron microscopy image of VLPSi. Scale bar is 50 nm. **(B)** Zeta potentials of VLPSi, amino-modified-VLPSi, and ALC-0315:DOPE (5:3 w/w) based transfection complexes. Significance: ***p ≤ 0.001.

**Figure 1b** presents the zeta potentials for both VLPSi with and without amino-modifications and transfection complexes. The zeta potential of VLPSi (-19.7 mV) was successfully turned to positive with all three amino-modifications, being 30.3, 27.4, and 34.0 mV for NH_2_-, N(CH_3_)_3_-, and NH-NH-NH_2_-VLPSi, respectively. After capping the free NH_2_ groups with COOH, the zeta potential of NH_2_-VLPSi-COOH was -19.3 mV. Moreover, the zeta potentials of -9.8 and -10.4 mV were determined for ALC-0315:DOPE (5:3, w/w) and -VLPSi, respectively. The zeta potentials of LNPs (-1.7 mV) and LNP-NH_2_-VLPSi (-4.1 mV) composed of ALC-0315, DOPE, and ALC-0159 (58.6 : 35.2 : 6.2, w/w) were close to neutral.

### 3.2 Cell uptake of VLPSi and LP-VLPSi

Our previous studies have indicated that spiky surface of VLPSi could enhance cell uptake as compared to the counterparts with spherical shape [21]. The rigid spikes on VLPSi possess piercing effect to disrupt cell membrane and thus, VLPSi is internalized mainly via caveolae-mediated endocytosis and micropinocytosis [29]. On the other hand, NPs prepared by lipids are mostly dependent on clathrin-mediated endocytosis and membrane fusion [30]. As shown in **Figure 2**, the uptake of both VLPSi and LP-VLPSi is minor during the first 1 h incubation. With the prolong of incubation time, more NPs are gradually internalized into cells. Notably, LP-VLPSi is more internalized into HEK-293T cells than VLPSi, indicating that there is a synergy between LPs and VLPSi for enhanced cell uptake.

**Figure 2.**
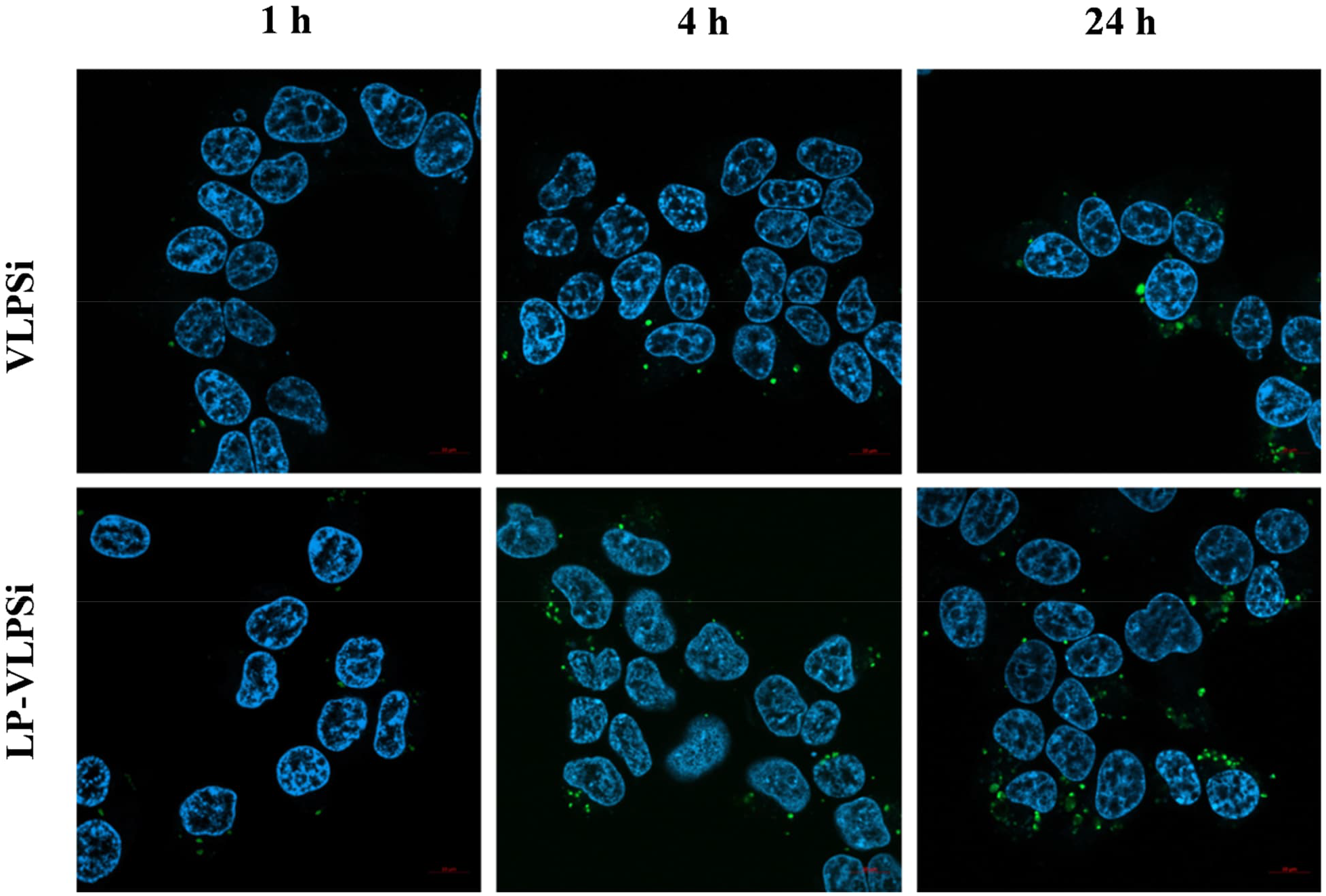
Fluorescence confocal images for studying cell uptake of VLPSi and LP-VLPSi into HEK-293T cells upon different incubation times. VLPSi was labelled with FITC (green), and cell nuclei were stained with Hoechst 33342 (blue).

### 3.3 mRNA transfection with LPs and LP-VLPSi

To study the effect of mass ratio between ionizable and helper lipid, ALC-0315 and DOPC were combined to form LPs using thin-film hydration method. **Figure 3a** shows the transfection efficiency of ALC-0315:DOPC with mass ratios of 1:1, 5:3, 2:1, 3:1, 4:1, and 5:1 in HEK-293T cells, when mRNA to LP mass ratio was 1:8. The highest 15% transfection efficiency was achieved with the mass ratio of 5:3, and it was selected for subsequent studies.

**Figure 3.**
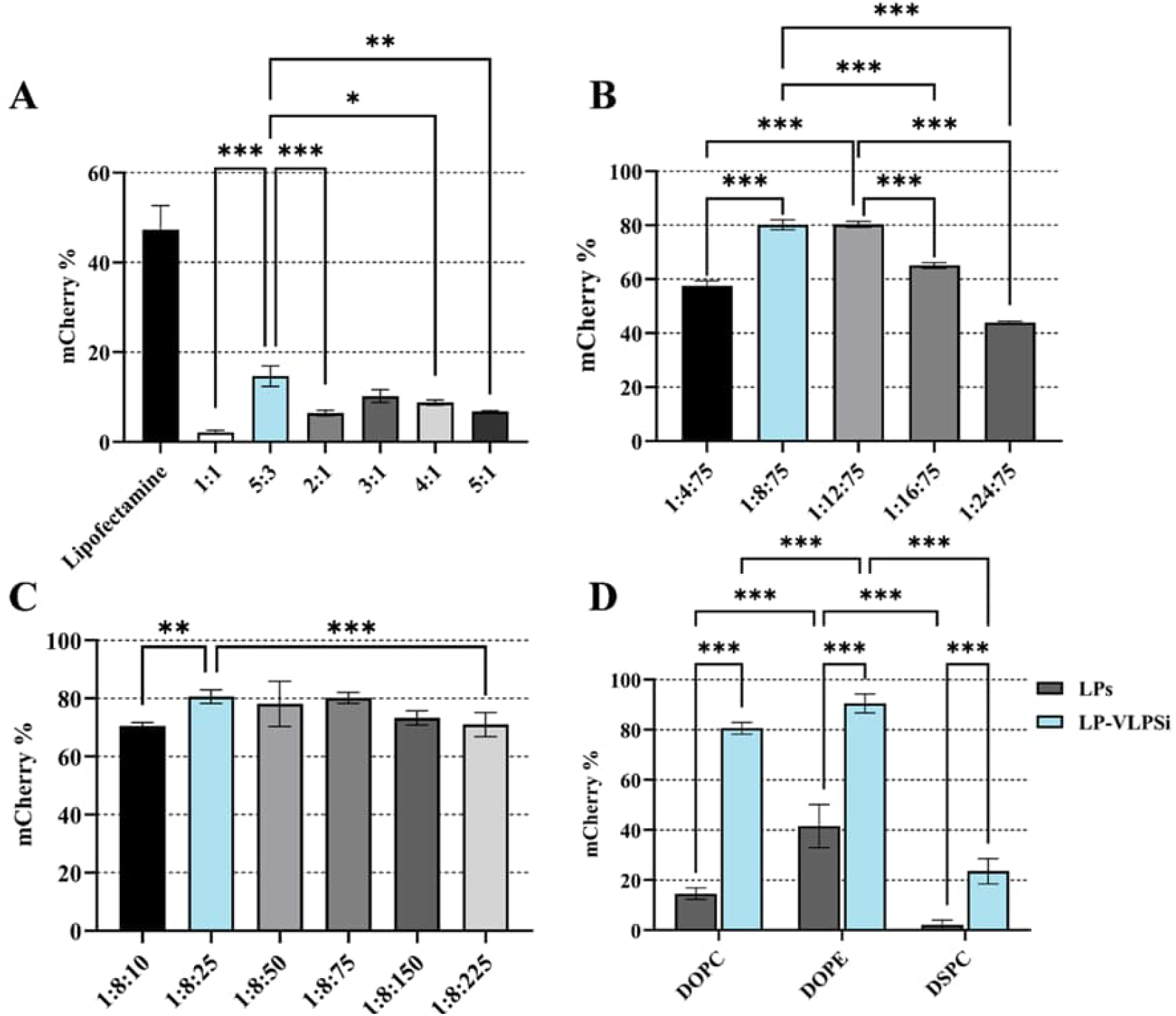
mCherry mRNA transfection with LPs and LP-VLPSi. **(A)** The transfection efficiency of Lipofectamine 2000 and ALC-0315:DOPC in the mass ratios of 1:1, 5:3, 2:1, 3:1, 4:1, and 5:1 in HEK-293T cells. The mRNA to LP mass ratio was 1:8. **(B-C)** The transfection efficiency of ALC-0315:DOPC (5:3) -VLPSi with different mass ratios between mRNA, LP, and VLPSi in HEK-293T. **(D)** The transfection efficiency of LPs and LP-VLPSi in HEK-293T cells, when LPs were composed of ALC-0315 and three different helper lipids, DOPC, DOPE, and DSPC in mass ratio of 5:3. The mass ratio between mRNA, LPs, and VLPSi was 1:8:25. Significance: *p ≤ 0.05, **p ≤ 0.01, and ***p ≤ 0.001.

Further, ALC-0315:DOPC (5:3, w/w) was combined with VLPSi. First, using HEK-293T cells, different LP amounts were tested while VLPSi mass was kept constant (**Figure 3b**). No significant difference between the mass ratios of 1:8:75 and 1:12:75 was observed. Both ratios achieved the mRNA transfection efficiency of 80%. Thus, further tests were continued with the lower mass ratio of 1:8 between mRNA and LPs, variating the amount of VLPSi (**Figure 3c**). The transfection efficiency reached 80% with the mass ratios of 1:8:25, 1:8:50, and 1:8:75. The result verified that the mass ratio of 1:8:25 with less VLPSi was sufficient for mRNA transfection.

In the presented transfection results (**Figures 3a-c**), DOPC was used as a helper lipid. Next, the effect of different helper lipids DOPC, DOPE, or DSPC was tested in the mass ratio of 5:3. The formed LPs were further combined with VLPSi in the mass ratio of 1:8:25. **Figure 3d** shows that the significantly highest mCherry expression in HEK-293T cells, with both LPs and LP-VLPSi, was obtained when DOPE was used a helper lipid. With DOPE, the transfection efficiency of LPs and LP-VLPSi was around 40% and 90%, respectively. In all the groups, LP-VLPSi achieved significantly higher mRNA transfection efficiency compared to LPs.

Based on the results presented in **Figure 3d**, DOPE was the most suitable helper lipid to form LP with ALC-0315. In the next test, two cationic lipids, DOTAP and DOTMA, were combined with DOPE in three different mass ratios: 1:2, 1:1, and 2:1. Subsequently, LPs were combined with VLPSi in the mass ratio of 1:8:25. No significant differences between different mass ratios of DOTAP and DOPE or DOTMA and DOPE was observed when transfection was studied with LPs in HEK-293T cells (**Figure 4a**). However, DOTAP:DOPE-VLPSi achieved significantly higher mCherry expression compared to corresponding LPs when the DOTAP to DOPE mass ratio of 2:1 was used. Moreover, significant difference in transfection efficiency between LPs and LP-VLPSi was observed in all three mass ratios of DOTMA and DOPE. The highest transfection of 60% was observed with DOTMA:DOPE (2:1) -VLPSi. However, the observed transfection was much lower than with ALC-0315:DOPE (5:3, w/w) presented in **Figure 3d**. Thus, ALC-0315:DOPE (5:3, w/w) was selected for the rest of the transfection experiments.

**Figure 4.**
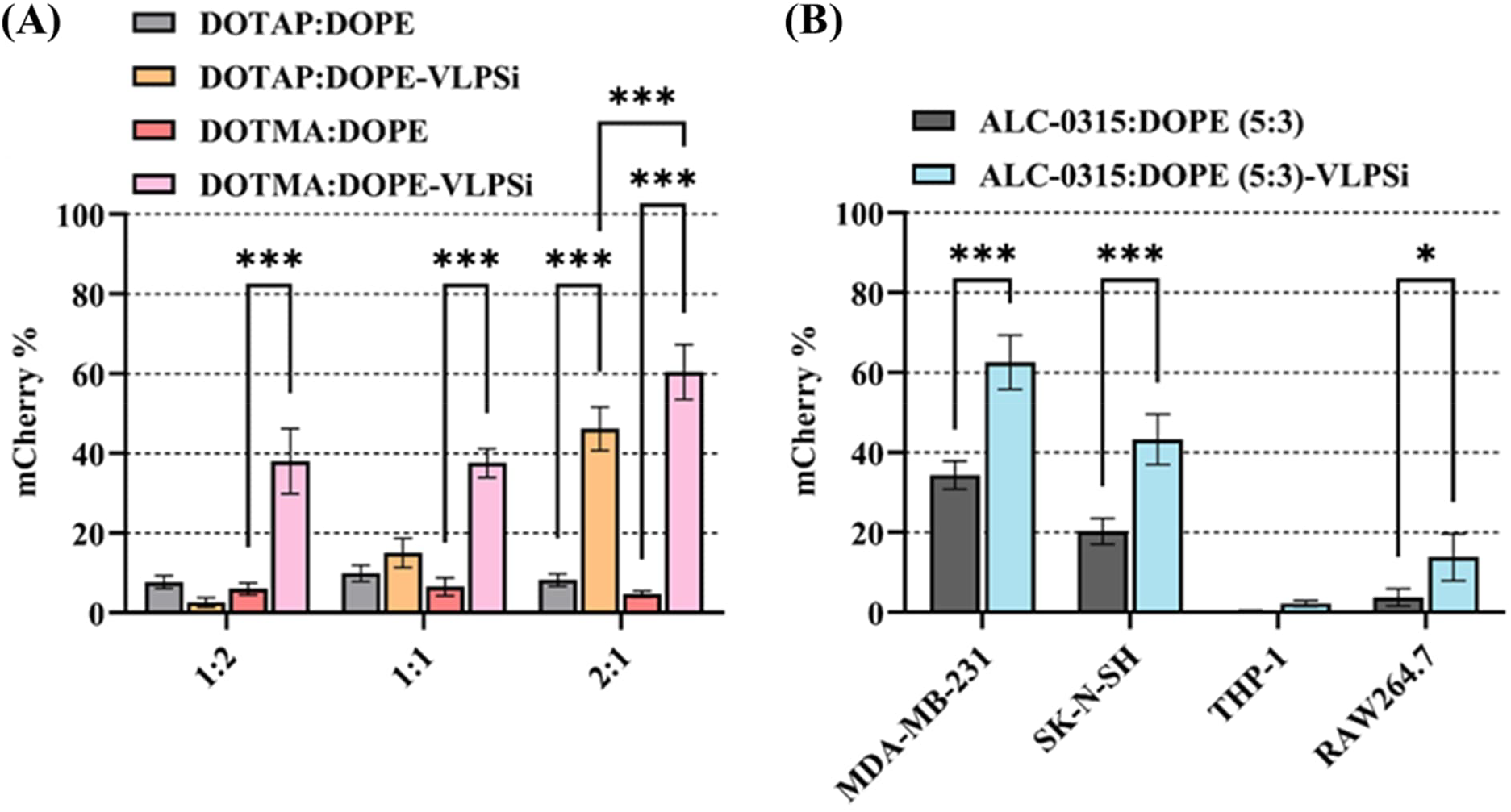
mCherry mRNA transfection with LPs and LP-VLPSi. **(A)** The transfection efficiency of LPs and LP-VLPSi in HEK-293T cells, when LPs were composed of DOTMA or DOTAP and DOPE in different mass ratios: 1:2, 1:1, and 2:1. The mass ratio between mRNA, LPs, and VLPSi was 1:8:25. **(B)** The transfection efficiency of ALC-0315:DOPE (5:3, w/w) with and without VLPSi in different cell lines. The mass ratio between mRNA, LPs, and VLPSi was 1:8:25. Significance: *p ≤ 0.05 and ***p ≤ 0.001.

HEK-293T cells were used to screen optimal lipid composition and mass ratios. To evaluate the transfection in other cell types, MDA-MB-231, SK-N-SH, THP-1, and RAW264.7 cells were tested with LPs and LP-VLPSi composed of ALC-0315 and DOPE (5:3, w/w). The mass ratio between mRNA and LPs and mRNA, LPs, and VLPSi were 1:8 and 1:8:25, respectively. The mRNA transfection efficiency variated between different cell lines as seen in **Figure 4b**. However, the transfection was significantly higher with LP-VLPSi compared to LPs in all other cell lines except THP-1. The highest transfection efficiency, around 60%, was achieved with LP-VLPSi in MDA-MB-231 cells.

### 3.4 mRNA transfection with LNPs and LNP-VLPSi

LNPs and LNP-VLPSi were prepared by pipette mixing method. LNPs were composed of ALC-0315, DOPE, and ALC-0159 in the mass ratio of 58.6 : 35.2 : 6.2. **Figure 5a** shows the mCherry expression in HEK-293T cells when LNPs and LNP-VLPSi with and without different amino modifications were used as transfection reagents. The mass ratios between mRNA and LNPs and mRNA, LNPs, and VLPSi were 1:8 and 1:8:25, respectively. The transfection efficiency of LNPs was significantly higher than LNP-VLPSi. However, when different amino modifications were applied on the surfaces of VLPSi, no significant difference in transfection efficiency between LNPs, LNP-NH_2_-VLPSi, and LNP-NH-NH-NH_2_-VLPSi was observed. All of these complexes achieved transfection efficiency of over 90%.

**Figure 5.**
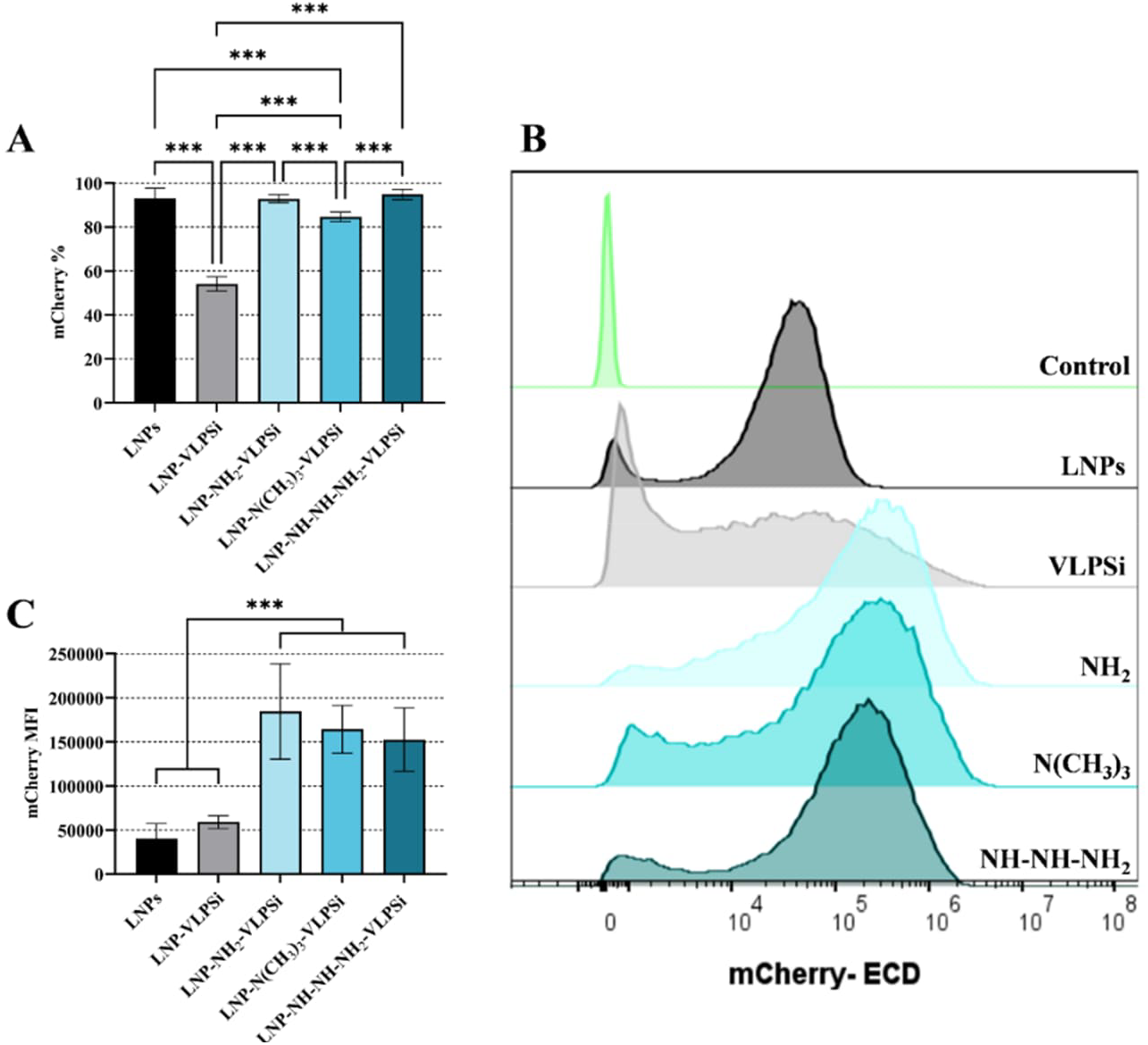
mCherry mRNA transfection with LNPs and LNP-VLPSi. **(A)** The transfection efficiency of LNPs and LNP-VLPSi with and without amino-modifications in HEK-293T cells. LNPs were composed of ALC-0315, DOPE, and ALC-0159 in the mass ratio of 58.6 : 35.2 : 6.2. The mass ratio between mRNA, LNPs, and VLPSi was 1:8:25. Corresponding **(B)** histograms of mCherry fluorescence intensity and **(C)** mCherry median fluorescence intensity (MFI) in HEK-293T cells. Significance: ***p ≤ 0.001.

To further assess mCherry expression, the median fluorescence intensity (MFI) was measured. **Figure 5b** shows histograms of mCherry MFI in HEK-293T cells for LNPs and LNP-VLPSi with and without different amino-modifications. By examining the histograms, a clear shift in mCherry intensity is visible for amino-modified VLPSi compared to LNPs or LNP-VLPSi. This is further confirmed in **Figure 5c**, showing the significant difference in mCherry MFI values between LNPs and amino-modified VLPSi. However, no significant differences between different amino modifications were observed.

## 4 Discussion

Biomimetic virus-like nanoparticles have aroused interest in several biomedical applications [20]. Recently, inspired by SARS-CoV-2 vaccines, the potential of VLPSi as an appropriate nuclei acid delivery system have been studied [24]. However, due to large size and negative charge of mRNA, its delivery efficacy has remained relatively low [25,26]. To enhance the mRNA delivery efficacy of VLPSi, some opposite charged linker between mRNA and VLPSi is needed. In the previous studies, MSNs have been coated with strongly cationic polyethylenimine via electrostatic binding for mRNA delivery [25,31]. Further, lipid bilayers have been introduced on the NP surfaces, supporting the mRNA binding [31]. However, LPs and LNPs containing positively charged cationic lipid or pH dependent ionizable lipid have proven their ability in mRNA delivery [28]. In those, the mRNA encapsulation is mainly based on electrostatic interaction, in which positively charged lipids self-assemble around negatively charged mRNA [27]. Thus, the idea of combining VLPSi with positively charged lipids to encapsulate mRNA for its delivery arose.

The findings of this study showed that both LPs and LNPs can be practically combined with VLPSi to obtain hybrid lipid-VLPSi for efficient mRNA delivery. The presented results confirmed that VLPSi can significantly increase mRNA transfection efficiency compared to both LPs and LNPs. This might be due to the viral spikes of NPs, which aid cell internalization and subsequently endosomal escape of mRNA by damaging endosomal membrane [23,24]. Further, the rigid VLPSi might improve the structural stability of mRNA complex and protect the mRNA during intracellular trafficking [32].

Previous studies have shown that different lipids and lipid compositions affect greatly to the mRNA delivery efficacy [33]. Especially, ionizable lipids have shown their promise over cationic lipids, becoming protonated in acidic endosomes [34]. This aims the destabilization of endosomal membranes and further release of mRNA into cytoplasm [15,34]. In this study, among the tested cationic and ionizable lipids, ALC-0315 was found to be the most effective for mRNA delivery in both LPs and LP-VLPSi. In addition, different helper lipids combined with ALC-0315 were tested, in which DOPE showed the highest mRNA transfection efficiency (**Figure 3d**). This could be explained by the ability of DOPE to further promote endosomal escape due to its destabilizing hexagonal structure in acidic environment [35].

Ideally, the positive surface charge of VLPSi could encapsulate mRNA via electrostatic interaction. In the previous study of Fu et al. (2022), they showed that amino-modification on VLPSi significantly promoted siRNA protection capability and delivery efficacy [24]. In our study, we demonstrated the importance of positive surface charge of VLPSi, when hybrid LNP-VLPSi were developed for mRNA delivery. Based on our findings, amino-modified VLPSi significantly increased the transfection efficiency compared to unmodified VLPSi (**Figure 5a**). Moreover, LNP-NH_2_-VLSPi achieved even five times higher gene expression than commonly used LNPs (**Figure 5c**), indicating the increased mRNA release from endosomes to cell cytoplasm.

As a summary, the present study introduced a novel hybrid lipid-VLPSi delivery system for efficient mRNA delivery. Among the tested LPs, ALC-0315:DOPE (5:3, w/w) was investigated as the most applicable lipid composition to be combined with VLPSi. Moreover, amino modification on VLPSi was found to be the crucial factor in mRNA delivery with LNP-VLPSi, expressing significantly higher mRNA intensity compared to conventional LNPs. Altogether, the straightforward integration of VLPSi into widely used LNP system was demonstrated, providing a flexible and efficient strategy for advancing next-generation mRNA therapeutics.

## Acknowledgement

This work was supported by the Research Council of Finland (Grant no. 356056). We acknowledge technical support from Petri Mäkinen from A.I Virtanen Institute for Molecular Sciences, and UEF Cell and the Tissue Imaging Unit, and SIB Labs.

