## Supplementary Information for "Bioinspired Virus-Like Porous Silica Amplify Lipid-Mediated mRNA Delivery"

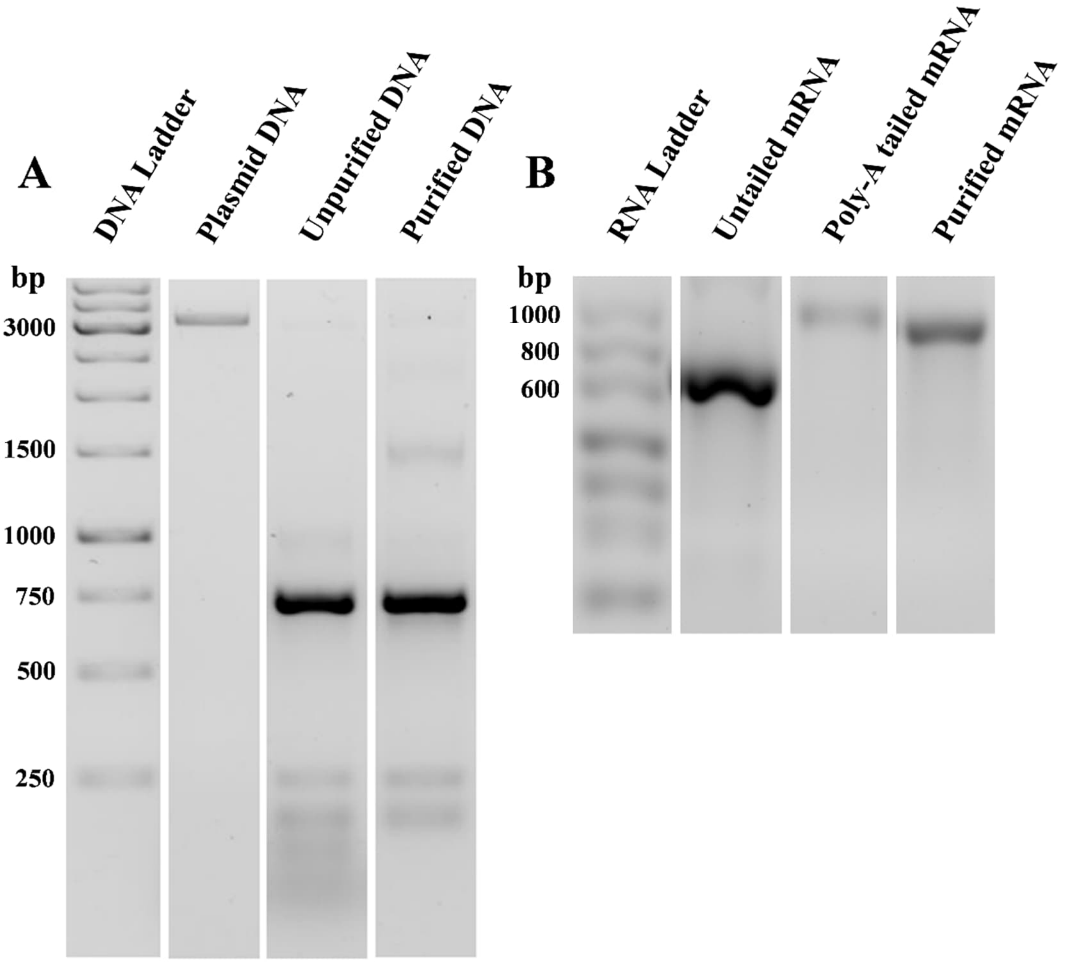


**Figure S1.** Confirmation of nuclei acid production and purity on agarose gel electrophoresis. **(A)** mCherry plasmid DNA and PCR amplified mCherry DNA before and after purification on 2% agarose gel containing 0.01% of GelRed. **(B)** mCherry mRNA before and after Poly-A tailing and final purified mCherry mRNA product on 1.5% agarose gels containing 0.01% of GelRed. The gels were imaged using ChemiDOC MP Imaging System (Bio-Rad, USA).

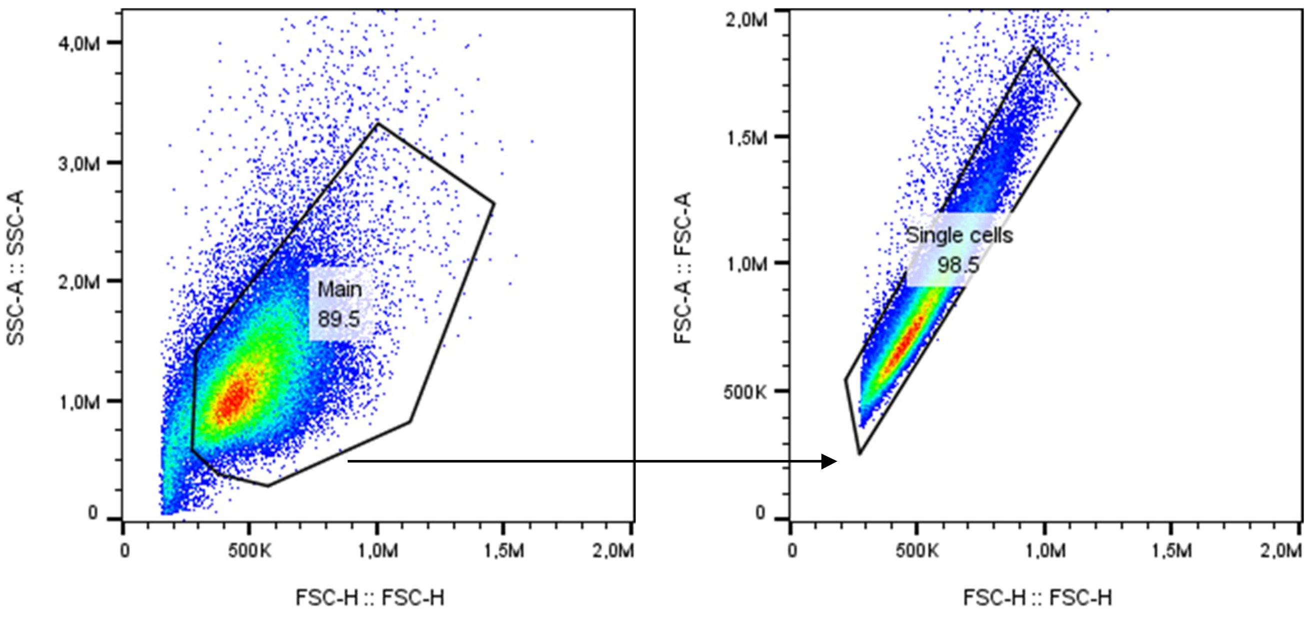


**Figure S2.** Flow cytometry gating strategy for the analysis of mCherry positive cells.
